# Pathological angiogenesis RNA network analysis reveals a VEGF-independent gene signature with prognostic power for HER2+ breast cancer

**DOI:** 10.64898/2025.12.16.694559

**Authors:** Jhonatas S. Monteiro, Lilian C. Alecrim, Ricardo J. Giordano, João C. Setubal

## Abstract

Cancer and many ophthalmic diseases share angiogenesis as a common element for disease progression. Using a well-accepted animal model, we performed network coexpression analysis of mRNA and regulatory RNA to investigate the molecular landscape of pathological angiogenesis in the mouse retina. We assessed the relevance of the RNA coexpression analysis by building gene signatures for breast cancer (METABRIC cohort). Most of the gene signatures were prognostic, and one was prognostic for HER2+ breast cancer, outperforming previously reported HER2+ signatures. Interestingly, it did not include *VEGF* as variable, and multivariate Cox analysis further highlighted *ANGPT2* as a key gene, suggesting that *VEGF*-independent neovascularization plays an important role in breast cancer. Next, we constructed a network linking mRNAs and regulatory RNAs involved in pathological angiogenesis; this analysis revealed three novel lncRNAs that may play a role in angiogenesis and tumor development. The novel prognostic gene signature and a comprehensive RNA-based regulatory network advance the understanding of angiogenic pathways and provide clinically relevant tools for HER2+ breast cancer stratification.

**Author Summary:** The formation of new blood vessels by angiogenesis is central in the origin and growth of cancer. Gene signatures are lists of genes that have strong correlation with a disease and/or its development. This study presents a novel angiogenesis gene signature with strong prognostic value for *HER2*+ breast cancer, an aggressive subtype of this disease. We also discovered novel long noncoding RNA genes that may have important roles in angiogenesis and in tumor development. These findings enhance our understanding of angiogenic mechanisms and offer clinically relevant tools for *HER2*+ breast cancer diagnosis, potentially guiding more precise therapeutic interventions.

## Introduction

Breast cancer is the most frequent cancer diagnosed in women in the world (although it also affects men to a lesser extent). At the molecular level, breast cancer is very heterogeneous, and it can be classified into four different subtypes based on its cell molecular expression profiles: luminal A and B (expressing the oestrogen receptor — *ER*), epidermal growth factor receptor (*HER2*) positive, and basal (*HER2*-negative and *ER*-negative) [1–3]. Although treatment options for most patients have advanced substantially in the past decades, breast cancer is still a disease with high mortality rate due to development of resistance to treatment, including anti-angiogenesis drugs. Angiogenesis is a hallmark of cancer, and anti-angiogenesis drugs have shown promising results for several types of cancer [4]. However, for breast cancer, anti-angiogenesis therapy has failed to improve patient survival, despite promising results in animal studies and initial approval from the FDA [5]. Thus, understanding the molecular behavior of metastatic breast cancer, including the identification of molecular markers and gene signatures with theranostic value is an important endeavor, which might contribute to improved diagnosis and treatment options to patients, including angiogenesis therapy.

Indeed, breast cancer treatment and diagnosis vary depending on tumor molecular subtype. For example, for many years the *HER2*+ subtype was associated with an aggressive phenotype and poor outcome, until the approval of *HER2*-targeted drugs, which brought a significant improvement in the survival outcome of these patients [6–8]. However, a portion of *HER2*+ patients, mainly those with tumors in the advanced stage (metastasis), will eventually develop resistance to existing therapies [6–8]. The mechanisms that lead to the resistance are not fully understood, thus, studying the signaling pathways and the genes involved in cancer progression and survival of *HER2*+ breast cancer may reveal new targets for the development of more effective therapies or novel prognostic markers to assist with treatment.

In previous work, we developed an angiogenesis-based gene signature that predicts breast cancer patient survival [9]. The gene signature was built using 153 differentially expressed genes (DEGs) in pathological angiogenesis, which were identified using a mouse model for retinopathy (oxygen-induced retinopathy — OIR). The OIR model is a powerful tool, extensively used to study pathological angiogenesis *in vivo* [10]. This is study was motivated by the well-known link between angiogenesis and cancer progression [4,11]. While, the gene signature worked well for the Luminal and Basal subtypes, it failed to classify patient with *HER2*+ breast cancer. Here, we revisited our work, added more data and performed further analysis in search of a gene signature that works well for the *HER2*+ subtype. For that, we used coexpression analysis, which is one of the methods that allows us to study relationships between different molecules that share the same biological function or pathway.

Hence, using WGCNA (Weighted Gene Co-expression Network Analysis) [12] to reanalyze the RNA-Seq data of the OIR mouse model [9], including unpublished results, along with new RNA sequencing data, we identified modules of coexpressed genes present in pathological angiogenesis. Next, we selected the best set of genes from those modules to build new gene signatures to identify those capable of distinguishing *HER2*+ breast cancer patients [9]. To further unveil the relationship between the molecules and their association with *HER2*+ breast cancer cells, we used the genes from the signature along with miRNA data from the OIR retina to build a mRNA-lncRNA-miRNA interaction network that revealed key regulators of the pathological angiogenesis pathway and that might be associated with breast cancer. As a result, these networks provided further insight into *HER2*+ breast cancer molecular behavior and expand the repertoire of candidates for the development of future therapies or prognostic markers for the treatment of breast cancer patients.

## Results

### Two hundred eighteen lncRNAs are differentially expressed in the angiogenic mouse retina, and 161 are new predictions

The RNA-seq datasets used in this study include data from our previous work [9], which comprised RNA-seq data from Poli-A+ mRNA, as well as data from a new RNA-seq dataset using total RNA depleted for ribosomal RNA (Ribominus) (The new data came from the same samples). In both cases, retina RNA was obtained from OIR mice models, a well-accept animal model for the study of pathological angiogenesis [13]. In brief, mice pups at postnatal day 7 (P7) were exposed to hyperoxia (75% O_2_ saturation) for 5 days and then returned to normoxia, creating a hypoxia condition in the retinas (Fig. 1A), while mice from the control group were kept under normoxia (20.8% oxygen condition) throughout the experiment. Retinas from both groups of animals were then dissected at P12, P12.5 (only for the OIR mice), P15, and P17 for RNA extraction and sequencing. From here on, samples from the OIR retinas will be referred to as R12, R12.5, R15 and R17, while samples from control mice will be referred to as P12, P15 and P17 (we did not collect retinas at P12.5) [9].

**Fig 1.**
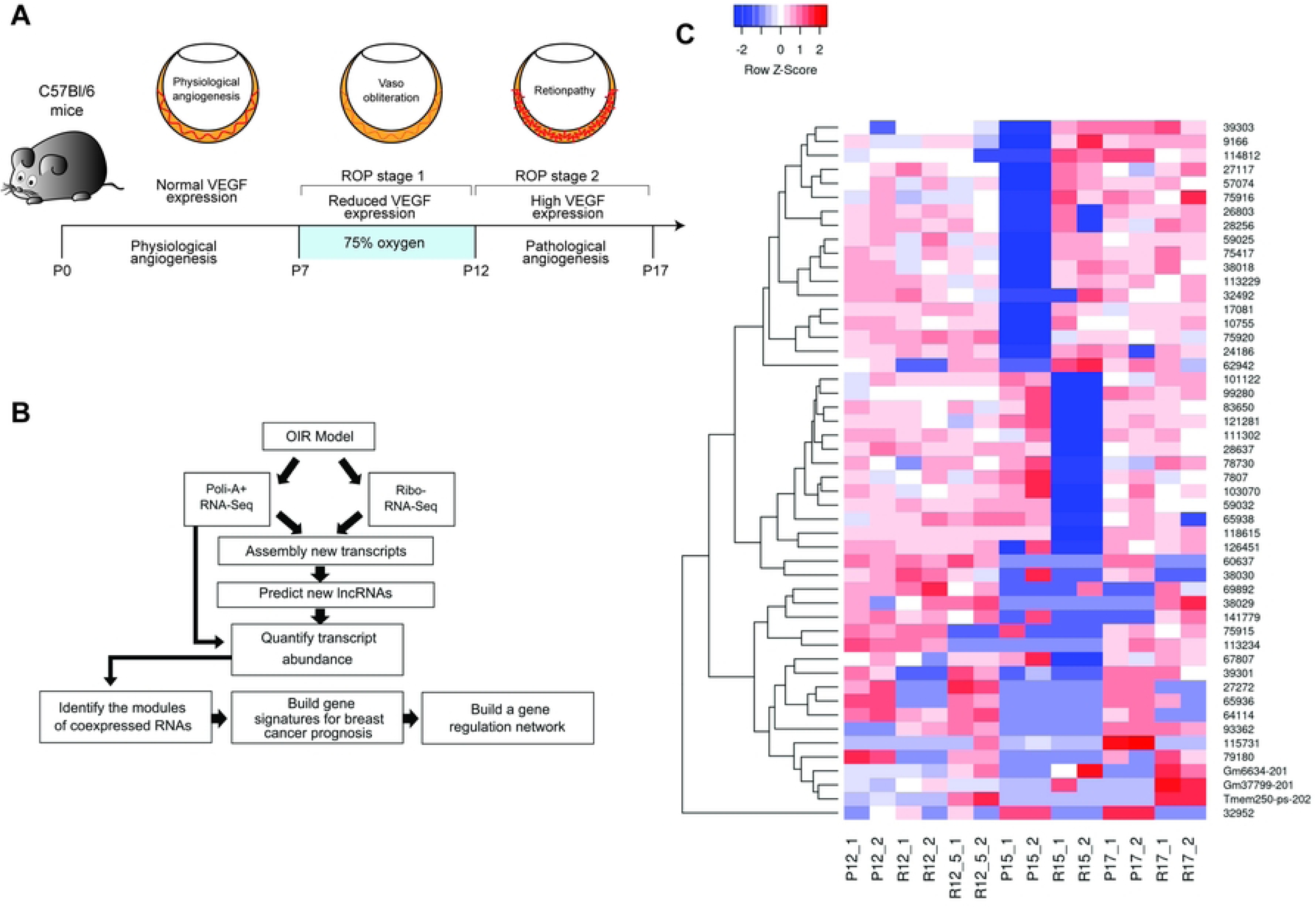
The OIR model and retina transcriptome reconstruction. (**A**) Illustration of the OIR model. Mice pups with their nursing mothers are kept in 75% oxygen for 5 days (P7 to P12), which reduces VEGF in mice retina leading to vaso-regression and central area vaso-obliteration. Upon return to room air, the now hypoxic retina increases VEGF levels resulting in a pathological angiogenesis state. (**B**) Overview of computational pipeline for the gene regulation network study. (**C**) Expression profile of 25 lncRNAs differentially expressed in pathological angiogenesis with the highest and lowest LFC (considering all possible permutations: R12 vs P12, R12.5 vs P12, R15 vs P15, and R17 vs P17. The letter P represents the samples from the control group, and the letter R represents the samples from the retinopathic group).

The RNA-seq reads from both datasets were combined and assembled, resulting in 52,926 transcripts, of which 10,492 were predicted to be lncRNAs based on their coding potential and similarity with other previously described lncRNAs (Table 1, Fig. 1B).

**Table 1.** Novel transcriptome assembly statistics.

| Statistics | # |
| --- | --- |
| Novel transcripts | 52,926 |
| Average size | 5,696 pb (min. 147 bp; max. 111.912 bp) |
| lncRNAs |  |
| CPAT + RNAsamba | 10,394 |
| BLAST | 181 |
| Infernal | 89 |
| Total new lncRNAs (non-redundant) | 10,492 |

Among the 10,492 lncRNAs, we observed that 218 were differentially expressed (DE) considering all time points: of those, 57 are previously known lncRNAs and 161 are new (Table 2, S1 Table). Most of the 218 DE lncRNAs were up-regulated during the angiogenesis phase (Table 2).

**Table 2.** Total of DE lncRNAs in each day (12, 15 and 17 postnatal days) of the experiment.

|  | R12 x P12 | R12.5 vs P12 | R15 vs P15 | R17 vs P17 |
| --- | --- | --- | --- | --- |
| Up-regulated | 2 | 26 | 48 | 110 |
| Down-regulated | 10 | 3 | 28 | 34 |

R, retinopathy group; P, control group.

The top 50 lncRNAs with the highest absolute log-fold change are 47 new ones and three previously described lncRNAs (*Gm6634-201, Gm37799-201*, and *Tmem250-os-202*) (Fig. 1C). The lncRNAs “62942” and “32952” had the highest and lowest values of LFC, respectively. lncRNAs “62942” is a non-coding isoform of the *MTPAP* gene and its expression varied throughout the angiogenesis process: following the hyperoxia phase (P12), it was 10-times lower in the retinopathic group, but its expression raised during the angiogenic phase and on P15 it was 10-times higher than the control group (S1A Fig.). On the other hand, the chromosomal coordinates of lncRNA “32952” indicate that it is a new isoform of the previously described lncRNA *MEG3*; its expression was reduced by up to 17 times in the retinopathic group on P17 (S1B Fig.).

### Modules of co-expressed genes in the mouse retina

The modules of co-expressed RNAs were inferred using WGCNA based on the normalized expression of the mRNAs and lncRNAs as input. Only RNAs with estimated variance above the median were considered in this analysis: 35,351 RNAs (25,218 mRNAs and 10,133 lncRNAs) out of 52,926 transcripts (66,7%). WGCNA identified 46 modules of co-expressed RNAs, which were ranked according to the number of transcripts they contained (mean = 751, max = 3,537 and min = 29) (Fig. 2A). By default, WGCNA labels modules using color names (Fig. 2B). The 29 transcripts that did not have any correlation, and therefore did not group, were included in the grey module (22 mRNAs and seven lncRNAs). A correlation analysis between the eigengenes of each module and the group of samples revealed that 23 modules (50%) had a significant correlation (*p*-value ≤ 0.05) with at least one group of the OIR model (Fig. 2C). Therefore, only these 23 modules were considered in downstream analyses.

**Fig 2.**
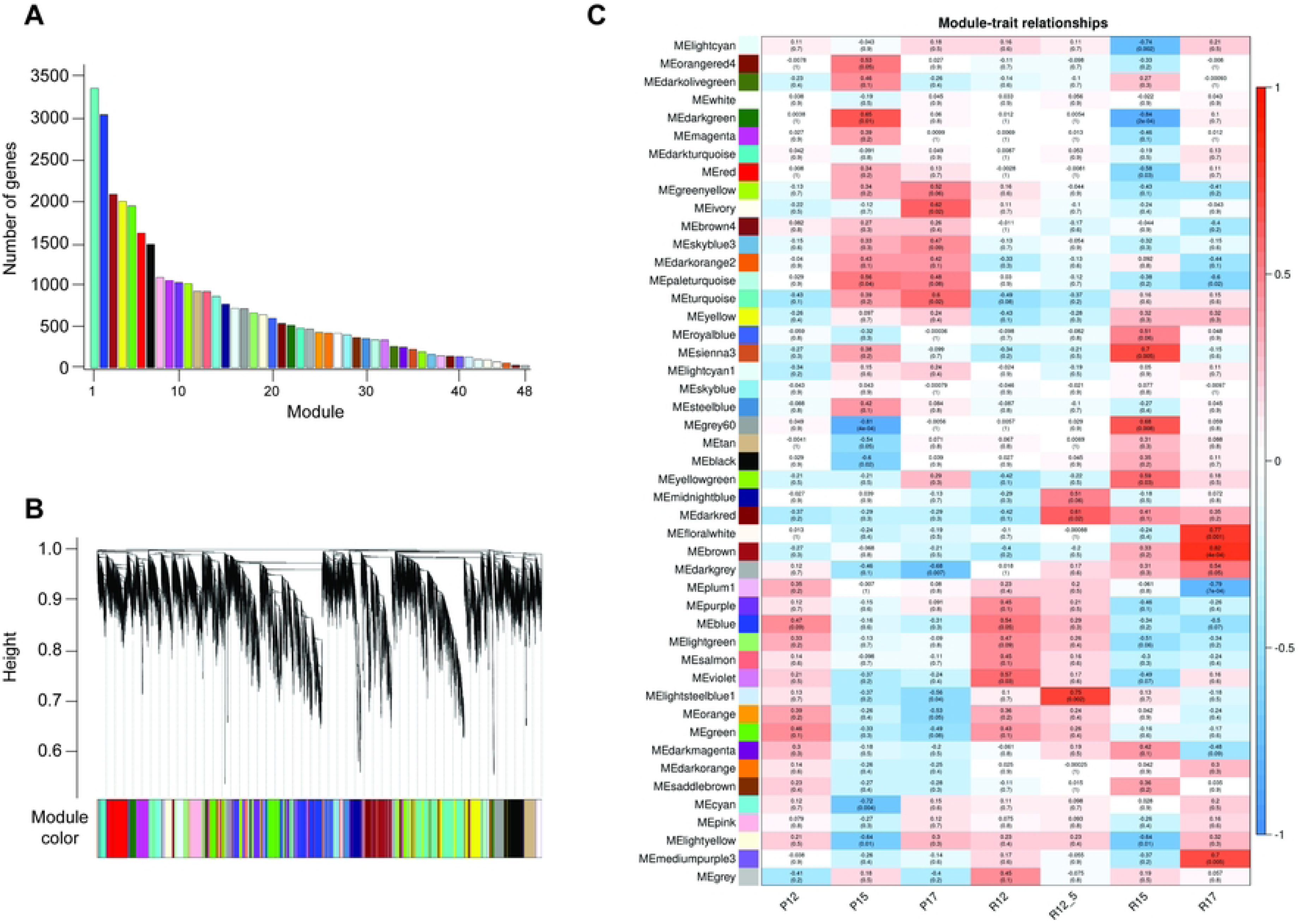
Construction of the angiogenic retina gene network. (**A**) Number of genes per module. (**B**) Dendrogram with the assigned colors to each gene network module. The genes were clustered using topological overlap measures[12] and the modules were defined using the Dynamic Tree Cut algorithm[115] as described in the experimental section. (**B**) Pearson correlation and the *p*-value for the eigengene of each module and the OIR condition. In total, there were 21 modules with a significant correlation with at least one group. The letter P represents the control group, while the pathological angiogenesis group is represented by the letter R, followed by the mice’s postnatal days (12, 12.5, 15 or 17).

### A novel and robust 44-gene signature for the HER2+ breast cancer subtype

Angiogenesis, a conserved mechanism present in vertebrate metazoans, has been proposed as an organizing principle to study angiogenesis-dependent diseases. Based on this idea, drugs and therapies for diseases as diverse as cancer and ophthalmic diseases have been developed [11].

As we did in our previous work, in order to assess the biological relevance of the WGCNA study we asked whether the identified genes would have prognostic value for angiogenesis-dependent diseases. In particular, given that our previous angiogenesis gene signature did not have prognostic value for the highly aggressive breast cancer *HER2*+ subtype, we focused our efforts on these particular patients. To develop a robust gene signature with prognostic power in *HER2*+ breast cancer, we employed the Random Forest model for high-dimensional feature selection and risk prediction, followed by Cox regression multivariate analysis to validate the signature’s statistical significance and quantify the independent contributions of individual genes, while adjusting for clinical covariates such as age and tumor stage. We again elected to use the dataset available at the molecular taxonomy of breast cancer international consortium (METABRIC) [14,15] for training and validation of two Random Forest models. METABRIC is one of the largest and most comprehensible datasets available for all types of cancer, as it contains gene expression profiles for 2,000 primary tumors that were split into two cohorts with no overlap: the training cohort (*n* = 961) and the validation cohort (*n* = 943) (Table 3). In what follows, to match the data in METABRIC, we only consider the mouse genes from the WGCNA modules that have human homologs.

**Table 3.** Clinical data of patients from the METABRIC database in the Training and Validation cohorts.

|  | **Training** | **Validation** |
| --- | --- | --- |
| Patients | 961 | 943 |
| Age at diagnosis (years)* | 61.26 (50.98, 70.22) | 62.61 (51.90, 70.93) |
| Overall survival (years)* | 9.70 (5.09, 15.43) | 9.46 (5.03, 15.26) |
| Tumor size* | 23 (18, 30) | 23 (17, 30) |
| **Tumor stage** |  |  |
| 0 | 1 | 3 |
| 1 | 300 | 175 |
| 2 | 517 | 283 |
| 3 | 72 | 43 |
| 4 | 8 | 1 |
| NA | 63 | 438 |
| **ER status** |  |  |
| Positive | 772 | 687 |
| Negative | 189 | 256 |
| **PAM50 subtype** |  |  |
| Basal | 69 | 130 |
| HER2 | 77 | 143 |
| LumA | 436 | 243 |
| LumB | 255 | 206 |
| Normal | 37 | 103 |
| NA | 0 | 6 |

NA = Not available; \* = Median (Lower Quartile, Upper Quartile).

Because the gene network modules contain hundreds of genes, we elected to build the gene signature with a maximum number of 50 genes, which were chosen based on the highest difference in expression between OIR and control retinas [16]. These 50 genes were then processed to remove genes that did not have a human homolog. After that, the remaining genes from each individual module were combined with patient age and tumor stage to feed the random forest package. The combination of genes with the lowest out-of-box (OOB) error was selected (minimum of four and maximum of 44 genes) (S2 Table): all 23 modules yielded gene signatures that could significantly segregate cancer patients (*log*-rank test *p*-value ≤ 0.05) using the METABRIC validation cohort into three categories of prognostic: low-risk (predicted probability of survival is equal to or higher than 50% in 15 years), intermediate-risk (between 7.5 and 15 years), and high-risk (less than 7.5 years) (S3 Table). These results are reassuring and validate our overall approach. However, when we consider the *HER2*+ breast cancer subtype alone, only the gene signature from the Brown (*log*-rank test *p*-value 2.03x10^-4^) and Ivory (*log*-rank test *p*-value 1.61x10^-2^) modules significantly segregated these patients (Fig. 3A and B). Even though the Brown and Ivory signatures were also able to segregate patients with different risk profiles for the LumA and LumB subtypes, both failed for the Basal subtype (S2 Fig.). This is likely due to the relatively small number of patients for the Basal subtype.

**Fig 3.**
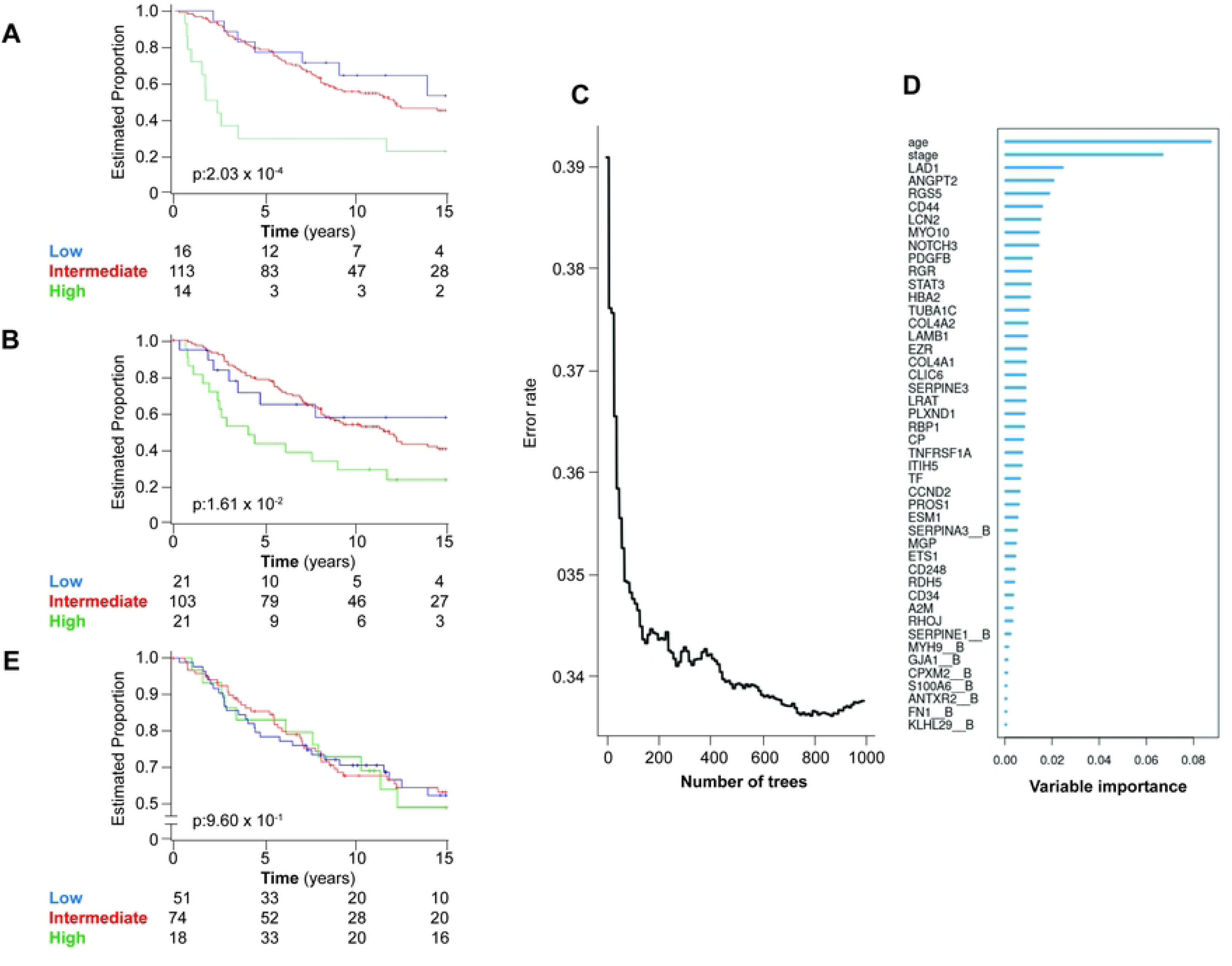
Gene signature for HER2+ breast cancer. (**A** and **B**) Kaplan-Meier curve for the Brown (A) and Ivory (B) gene signature in HER2+ breast cancer subtype of the METABRIC cohort. (**C**) The cumulative out-of-bag (OOB) error rates for each tree grown by the random forest algorithm using the Brown gene signature as a model. (**D**) Variable importance (VIMP) of each feature in the Brown gene signature. The suffix “ B” represents the genes with dichotomized expression values, where 0 is when the gene expression is below the median (considering the expression values in all samples) and 1 when is above the median. (**E**) Kaplan-Meier curve for a gene signature using only age and cancer stage and HER2+ breast cancer subtype of the METABRIC cohort. (**A, B and E**): *p*-value was calculated using log-rank test. Patients were classified in one of the risk categories: low-risk, if the predicted probability of survival is equal or higher to 50% in 15 years, intermediate-risk if between 7.5 and 15 years, and high risk if less than 7.5 years.

On closer inspection, we observed that the gene signature derived from the Brown module had a much stronger statistical power (*p*-value = 0.000203) than the gene signature from the Ivory module (*p*-value = 0.016) and that genes within the Brown module had the highest correlation (*p*-value = 0.00037) with genes expressed at the peak of the pathological angiogenesis process (R17 samples), compared to genes in the Ivory module (*p*-value = 0.019). Furthermore, only the genes in the Brown module showed gene ontology enrichment in functional analysis, while the genes in the Ivory module showed no such enrichment (S3 Table). Therefore, we decided to continue our analysis only with the gene signature derived from the Brown module, focusing on *HER2*+ patients.

The Brown module signature has 46 features (tumor stage + age + 44 genes with human homolog co-expressed in OIR mouse retinas) and achieved an accuracy of 66.2% (Fig. 3C). To further evaluate the performance of our gene signature, we performed two additional analyses. First, we asked whether random gene signatures could perform similarly. Using only genes significantly associated with the overall survival (OS) of breast cancer patients (Cox regression univariate analysis *p*-value ≤ 0.05) (S1 File), we built 10,000 signatures containing between 40 to 50 genes chosen at random (with replacement) in order to verify whether the specific gene composition from the Brown module actually contributes to the predictive power of the signatures. We observed that the great majority (97.5%) of all the gene signatures failed to predict the outcome for *HER2*+ patients. Moreover, our model had a more significant log-rank *p*-value than 99.91% of all random signatures.

Next, we analyzed the contribution of the two non-gene features: patient age and tumor stage. These two variables are of the highest importance to the signature (Fig. 3D). However, alone they fail to segregate *HER2*+ patients (Fig. 3E), clearly indicating that the genes from the Brown module are essential for its prognostic power. Taken together, these results reinforce our overall approach with WGCNA and the contribution of regulatory RNAs to the identification of genes with prognostic power to predict the survival for patients with *HER2*+ breast cancer subtype.

In order to identify which genes are the most robust individual predictors in the signature after adjusting for all other variables, we performed a multivariate Cox regression analysis. While the Random Forest model handles high-dimensional data well by considering interactions/nonlinearities, Cox regression tests each gene’s independent linear contribution while adjusting for others factors. Thus, the 44 genes with human homologs from the Brown signature were next selected as covariates to feed to the multivariate Cox regression analysis using the METABRIC validation cohort. This analysis again displayed a strong correlation and prognostic performance, confirming that the gene signature as a whole is strongly associated with survival in the validation cohort (Table 4).

**Table 4.**
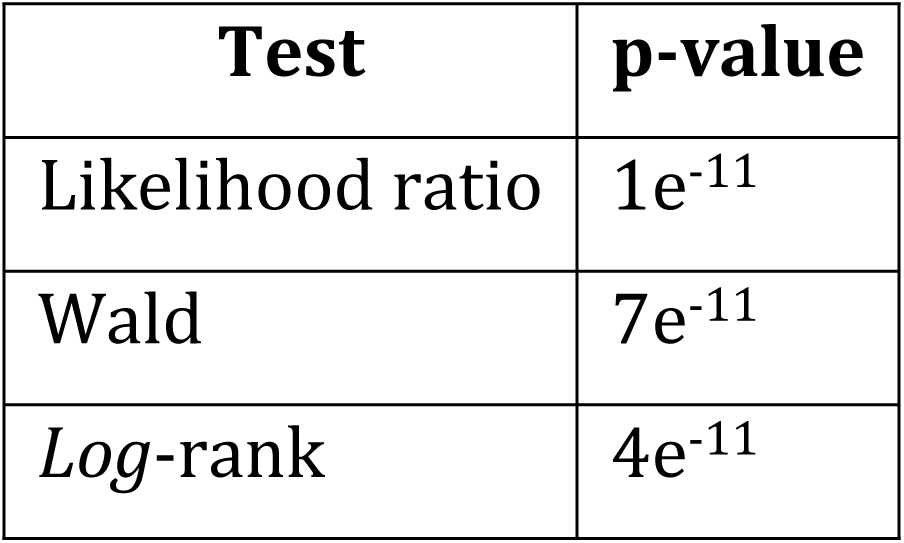
Cox regression multivariate analysis results using genes from Brown signature as covariates and the validation cohort as dataset.

Even though most of the 44 genes alone as covariates did not show a significant correlation with patient outcome, we observed that five genes had significant estimates for hazard ratios (HR) as covariates affecting survival (Fig. 4): *CCND2 and CLIC6* with HR < 1 predicts a better survival (protective gene), while *TUBA1C*, *ANGPT2* and *EZR* with HR > 1 predict worse survival (risk genes), all of which have already been associated with breast cancer [17–22]. Finally, to understand the role of the genes found in the Brown module and their possible contribution to breast cancer progression, we performed enrichment function analysis with Gene Ontology (GO) [23]. It revealed that genes from the Brown module signature are related to the regulation of angiogenesis and vascular development, and peptidase activity; more specifically, inhibition/regulation of peptidase activity (Fig. 5).

**Fig 4.**
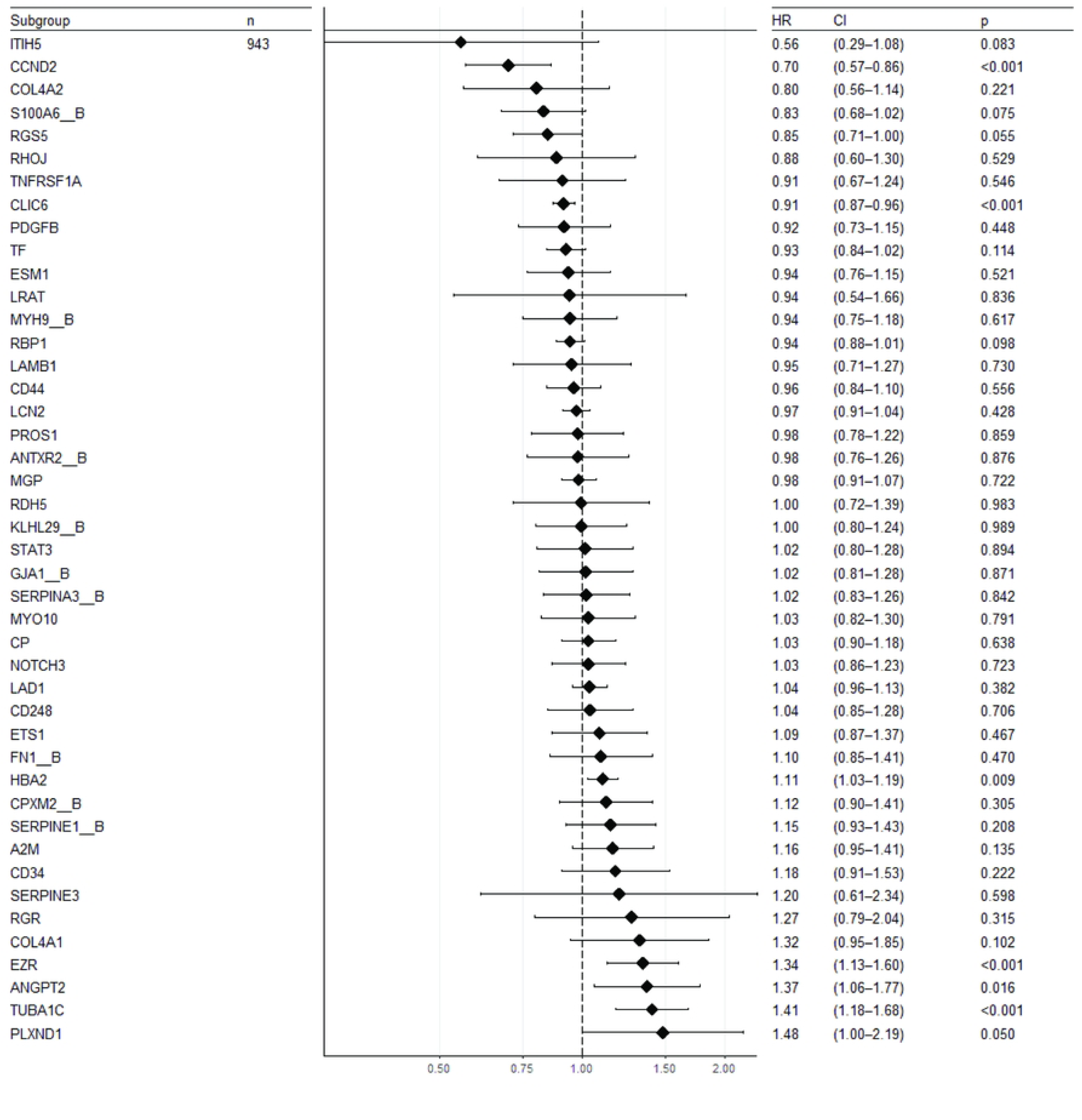
Genes with significant correlation with patient outcome. The plot shows the hazard ratio (HR), confidence interval (Cl) and significance (*p*-value) for each gene from Brown signature used as covariates in a Cox regression multivariate analysis using the METABRIC validation cohort as dataset.

**Fig 5.**
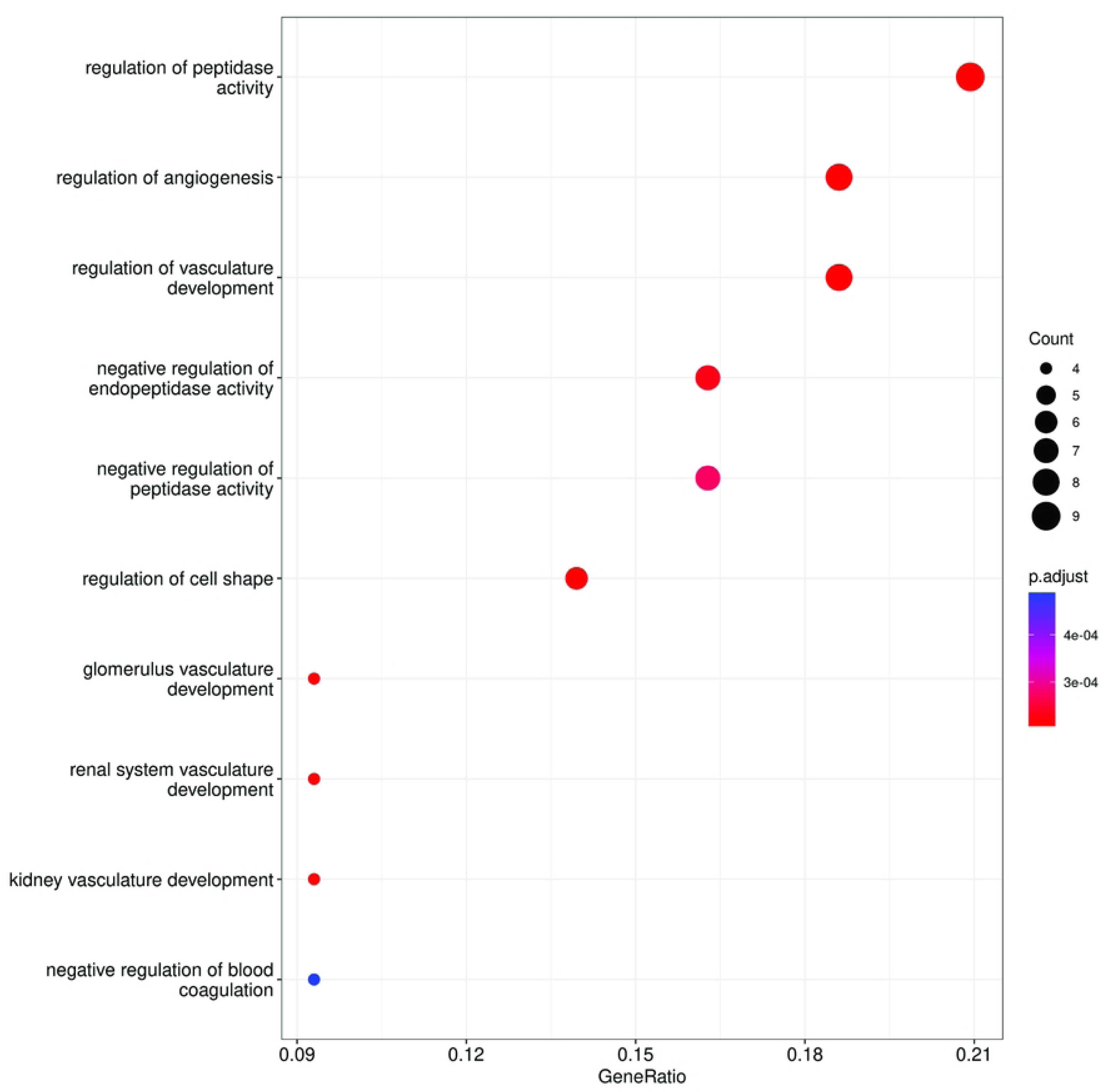
Brown signature gene ontology. The top 10 GO terms from the Biological Process ontology that were enriched in the Brown genes signature. Circle diameter indicate number of genes associated with each individual biological process and circle color the *p*-value.

### The new signature compares favorably against previously published HER2+ gene signatures

Having validated our gene signature for *HER2*+ breast cancer, we compared it with other four gene signatures published previously [6,24–26]. In all cases, the specified set of genes from each signature plus patient age and tumor stage was used to feed the Random Forest model using the METABRIC dataset in the same way the Brown module signature was built. The results indicate that the Brown module gene signature was superior to all other signatures, with only one of the published signatures displaying a prognosis similar to the Brown module signature (based on *log*-rank *p*-value) (Table 5).

**Table 5.**
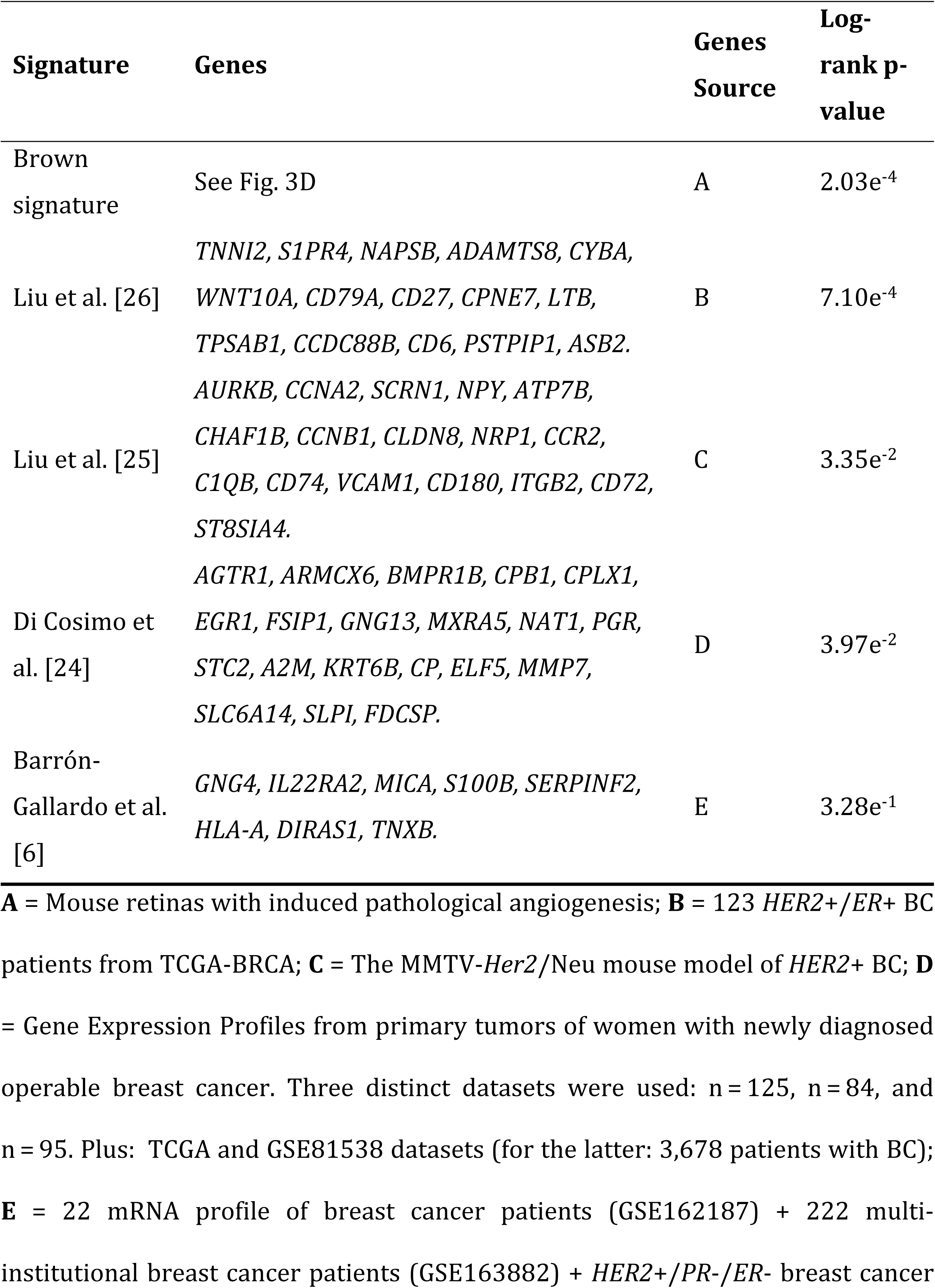
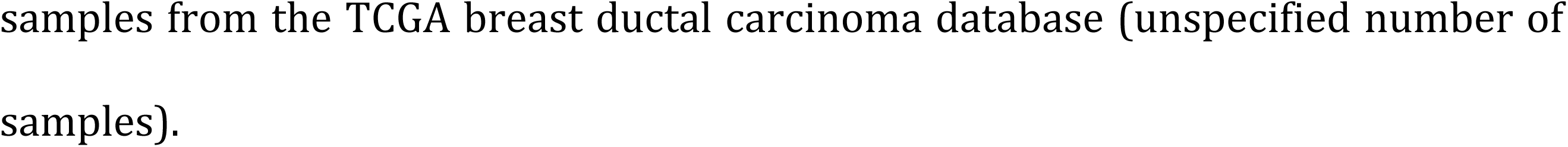
A comparison of the predictive power of the Her2+ gene signatures previously published in the literature and the brown gene signature using the gene expression data and clinical data (tumor stage and age of the patient) of the *HER2+* patients from METABRIC validation cohort.

Interestingly, the five signatures have almost no genes in common, with the notable exception of two genes, *A2M* and *CP*, shared by the Brown module signature and the Di Cosimo et al. [24] gene signature (Fig. 6). *A2M* encodes the alpha-2-microglobulin, a protease inhibitor and cytokine transporter (corroborating our GO analysis about the importance of endopeptidase regulation), while *CP* encodes ceruloplasmin, a cupper and iron transport protein.

**Fig 6.**
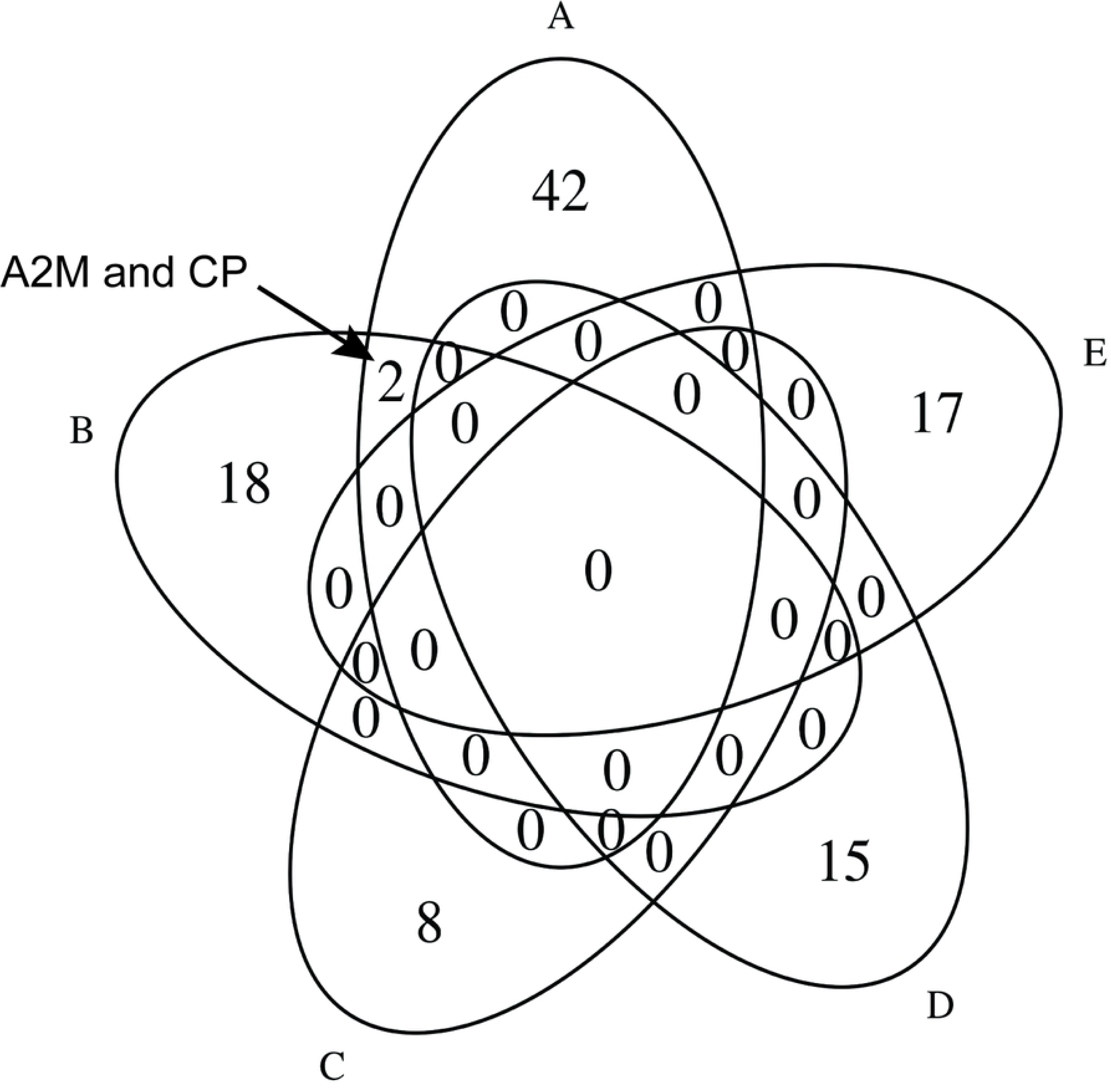
Genes shared among distinct breast cancer HER2+ gene signatures. Venn diagram comparing the set of genes of each signature. A = Brown signature, B = Di Cosimo et al.[24], C = Barrón-Gallardo et al.[6], D = Liu et al.[26] and E = Liu et al.[25].

In sum, the Brown module signature is a novel gene signature with better performance than previously published signatures, with potential applications in clinical settings.

### Two newly predicted lncRNAs seem to be important regulators in breast cancer

In the next step, we sought to gain additional insights into the molecular interactions associated with *HER2*+ and angiogenesis. For that, we used STRING [27], a database of known and predicted protein-protein interactions, to retrieve a protein-protein interaction (PPI) network. Only 20 of 44 genes from the Brown module signature had PPI data available in the STRING database. For the other 24 genes without PPI data in STRING and the lncRNAs of the Brown module, we postulated mRNA-mRNA or lncRNA-mRNA interactions for those pairs with WGCNA interaction weights above 0.2519518 (90^th^ percentile of all interaction weights of the module) (Fig. 7A); we obtained the final regulatory network by merging these two types of interactions.

**Fig 7.**
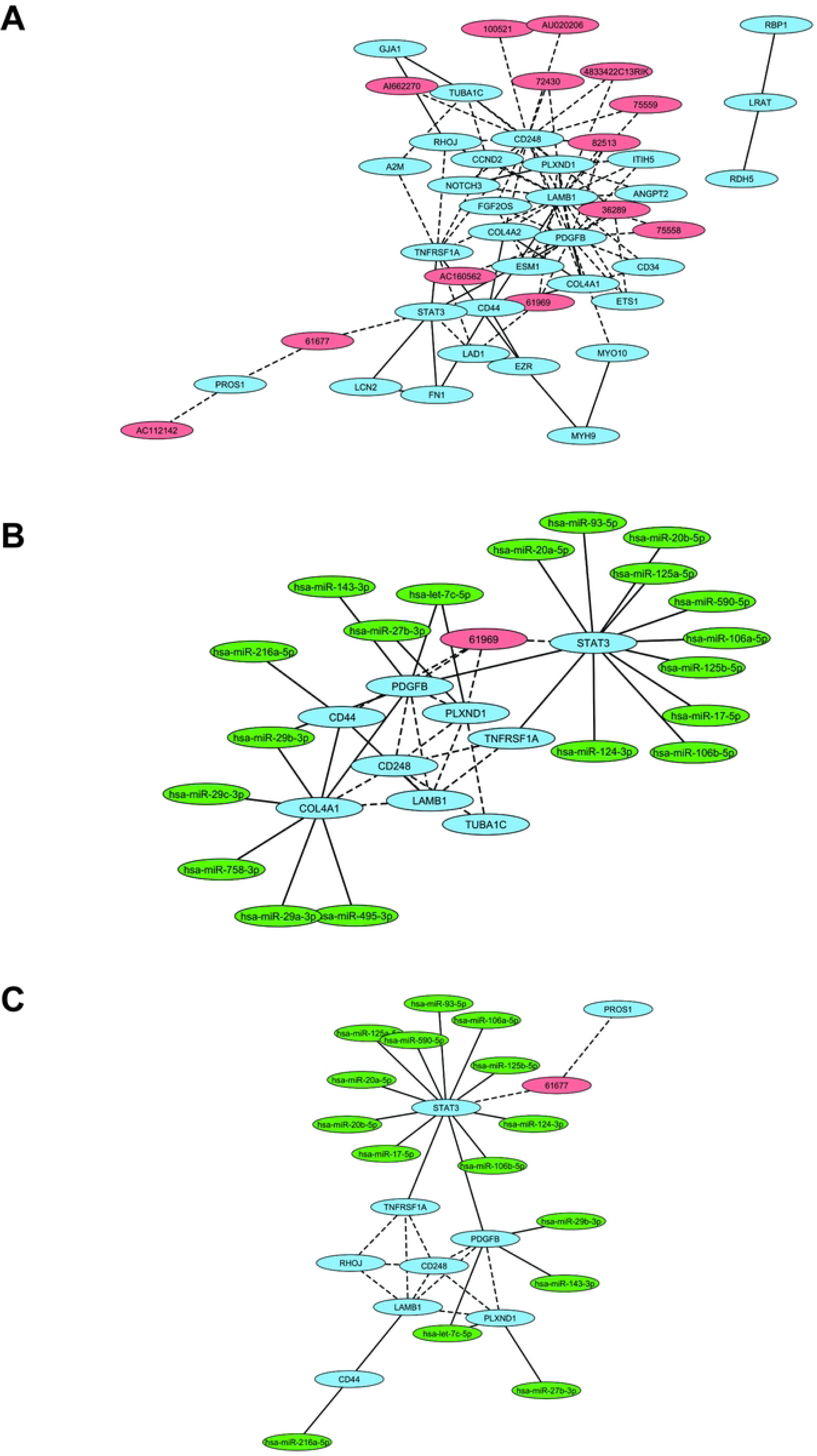
Gene network between retinal mRNAs and regulatory RNAs. (**A**) Interaction network of mRNAs and lncRNAs involved in regulating pathological angiogenesis. Solid edges represent protein-protein interactions found in the STRING database, while dashed edges represent the mRNAs or lncRNAs that are co-expressed in mouse retinas. (**B**) Subnetworks with the top 10 nodes with the highest degree in the pathological angiogenesis regulation network (**A**), and their potential MTI (miRNA:Target Interaction). (**C**) Subnetwork with the top 10 nodes with the highest betweenness centrality in the pathological angiogenesis regulation network, and their potential MTIs. (**B, C**) Solid edges represent high-confidence interaction (STRING and miRTarBase results), while dashed edges represent low-confidence interaction since the only evidence for these interactions is that they are co-expressed in mouse retinas. Red ellipses are lncRNAs, blue ellipses are the mRNAs, and the green ellipses are the miRNAs.

Using Cytoscape [28] to calculate the degree and the betweenness centrality of each node in this network, we ranked the nodes based on these two statistics (Table 6).

**Table 6.** Top 10 nodes with the highest degree and betweenness in the pathological angiogenesis regulation network.

| Rank | Degree | Rank | Betweenness |
| --- | --- | --- | --- |
| 1<sup>o</sup> | *CD248* | 1<sup>o</sup> | *PDGFB* |
| 2<sup>o</sup> | *TUBA1C* | 2<sup>o</sup> | *CD248* |
| 3<sup>o</sup> | *COL4A1* | 3<sup>o</sup> | *LAMB1* |
| 4<sup>o</sup> | *LAMB1* | 4<sup>o</sup> | *STAT3* |
| 5<sup>o</sup> | *CD44* | 5<sup>o</sup> | *TNFRSF1A* |
| 6<sup>o</sup> | *TNFRSF1A* | 6<sup>o</sup> | *61677* |
| 7<sup>o</sup> | *61969* | 7<sup>o</sup> | *PROS1* |
| 8<sup>o</sup> | *PLXND1* | 8<sup>o</sup> | *CD44* |
| 9<sup>o</sup> | *PDGFB* | 9<sup>o</sup> | *PLXND1* |
| 10<sup>o</sup> | *STAT3* | 10<sup>o</sup> | *RHOJ* |

The only lncRNAs in both lists are new lncRNAs predicted in our pipeline: “61969” (one of the top 10 nodes with the highest degree) and “61677” (one of the top 10 nodes with the highest betweenness centrality). Both lncRNAs were upregulated on days 15 and 17, and only lncRNA “61677” was upregulated on day 12 (during the hypoxia peak) (S1 Table). lncRNA “61969” is on the opposite strand of gene *RAB12*, which is in the green module (S3A Fig.). lncRNA “61677” is on the opposite strand of gene *RFX2* (S3B Fig.), which is also in the brown module, meaning that the mRNA from gene *RFX2* and lncRNA “61677” are co-expressed (interaction weight of *RFX2-61677* was 0.23144255, very close to the minimum threshold used to build the network). Interestingly, the *RFX2* gene encodes a transcription factor that regulates the expression of several oncogenes and cancer suppressor genes, and has been proposed as a promising therapeutic target for cancer treatment [29]. It is often mutated in breast cancer [29].

All genes associated with the top 10 nodes with the highest degree and betweenness in the pathological angiogenesis regulation network (Table 6) have been associated with angiogenesis. For this reason, we next looked for miRNAs that have the potential to regulate the expression of these genes. We found 19 potential miRNAs targeting the central genes of the pathological angiogenesis regulation network (Figs. 7B and C). These inferred networks of regulatory RNAs should be viewed with caution. Nevertheless, the observed correlations offer partial proof-of-concept to warrant further validation of these individual genes and regulatory RNAs and their participation in breast cancer.

## Discussion

Despite promising results in animal models, anti-angiogenic therapies have failed to improve OS in metastatic breast cancer. The reasons for this are yet unknown, and different hypotheses have been proposed, such as angiogenesis mediated by growth factors other than *VEGF* (i.e., *FGF2*, *ANG2*) (given that most anti-angiogenesis drugs are also anti-*VEGF*) and vessel co-option, among others [30,31]. Indeed, in our previous work, we showed that genes up-regulated in angiogenesis could predict the survival of breast cancer patients. Yet, our gene signature failed to provide a prognostic for breast cancer patients with the *HER2*+ subtype, which is a more aggressive form of breast cancer.

Here, using a larger dataset of angiogenesis-associated RNAs and supervised and unsupervised machine learning techniques, we developed a new 46-feature signature capable of predicting OS of patients with *HER2*+ breast cancer and classifying them into three categories of risk. It is important to emphasize that this new signature not only works on the *HER*2+ cancer subtype, but also for the LumA and LumB subtypes, and also for a mix of these subtypes (S3 Table), which makes this signature useful to a variety of patients. The genes from our signature were derived from the brown module of co-expression, and are functionally enriched in the regulation of angiogenesis, including a novel feature: peptidase activity. Peptide activity is associated with a broad number of biological processes, but it also features prominently in cancer and vascular cell migration associated with metastases and vessel formation, as well as extracellular matrix remodeling. Notably, the *HER2*+ gene signature lacks *VEGF*, which was present as a prominent feature in a previous angiogenesis signature [9], suggesting that neovascularization in *HER2*+ breast cancer might be driven by other pathways, such as *ANGPT2* (an important feature in our new signature).

As we did in our previous study, we compared our novel signature with other published gene signatures for *HER2*+ breast cancer. Although we found only four other gene signatures, and all of them were distinct regarding the genes they contained, ours ranked higher and more significantly (*p*-value) when classifying *HER2*+ patients (albeit somewhat close in *p*-value to the signature of Liu and collaborators [26]). Interestingly, with two exceptions, none of the gene signatures have genes in common. The notable exception was the signature by Di Cosimo et al. [24] and ours, which have two genes in common: *A2M* and *CP*. Alpha2-macroglobulin (*A2M*) is a pan-proteinase inhibitor whose overexpression in cancer cells is related to a better prognosis [32,33]. Low-risk *HER2*+ patients expressed more *A2M* compared to intermediate risk patients (*p*-value = 0.012) (S4 Fig.), agreeing with previous studies [32,33]. Ceruloplasmin (*CP*) is a ferroxidase mainly synthesized in the liver and secreted to the circulatory system, and it is responsible for the regulation of iron and copper levels[34,35]. *CP* has been found overexpressed in different cancer cells [35–40], including breast cancer [34].

Considering that anti-*VEGF* drugs have failed to improve breast cancer patient OS, it is conceivable that genes from the *HER2*+ signature might be possible targets for new anti-angiogenesis therapies towards cancer treatments. For instance, both the *ANGTP2* and *PDGFB* (platelet derived growth factor subunit B) genes are features in our *HER2*+ signature and encode growth factors that are proangiogenic and responsible for the recruitment of perivascular cells to augment formation and stabilization of new vessels [41]. Their expressions are associated with tumor progression in different types of cancer [41–45]. Furthermore, *PDGFB* overexpression has been associated with poor outcome for *HER2*+ breast cancer patients [46]. In our study, *ANGPT2* predicts worse survival (HR = 1.37, *p*-value = 0.016). Interestingly, *CD248* (endosialin/tumor endothelial marker 1) is also featured in the gene signature and encodes another proangiogenic factor that is expressed during tumorigenesis and embryogenesis, but not in adult tissues [47–49], which makes it an ideal target for the development of drugs for the treatment of cancer and other angiogenesis-dependent diseases. *CD248* acts together with the *PDGF* signaling pathway in the proliferation of pericytes during angiogenesis. Blockage of *CD248* leads to inhibition of pericyte proliferation induced by *PDGFB* [48]. *CD248* was coexpressed with *PDGFB* in the OIR model. Another gene associated with *PDGFB* pathways is *CD44*, which encodes a transmembrane glycoprotein expressed in most cell types, and a proangiogenic factor that mediates pathological angiogenesis [50] as well as tumor progression and metastasis in a variety of cancers [50,51], including breast cancer. In breast cancer it has been observed that *CD44* activates the *PDFGR-β*/*STAT3* signaling pathway to promote breast cancer stem cells. In agreement with these observations, in our analysis *CD44* was seen to be co-expressed with *PDGF-B* and *STAT3*.

In terms of degree and betweenness, the top 11 genes in the regulatory network (Fig. 7) are *TUBA1C, PROS1, STAT3, PDGFB, CD44, PLXND1, TNFRSF1A, CD248, COL4A1, LAMB1,* and *RHOJ. PDFGB, CD44* and *CD248* were discussed above. We now discuss the others.

*TUBA1C* is a subtype of α-tubulin that constitutes microtubules [52–54] and is primarily involved in the dynamic process of polymerization and depolymerization by cell division and replication [52,54]. Its overexpression has commonly been associated with poor prognosis for lung [55], liver [53,56], pancreas [52], brain, and spine cancer [54]. Our analysis also associated *TUBA1C* expression with a poor outcome for breast cancer patients (HR 1.41, *p*-value < 0.001) and *TUBA1C* also features in the regulatory RNA network associated with gene responsible matrix remodeling, and cell migration and survival. For instance, *TNFRSF1A* is a transmembrane receptor of the tumor necrosis factor-α (*TNF-α*). When it is activated it promotes tumor growth, invasion and metastasis in breast cancer cell lines by activating the NF-κβ signaling pathway [57–59] and promotes gastric tumorigenesis by inducing *NOXO1* and *GNA14* genes [60]. *TNFRSF1A* has a binding site for the *STAT3* gene, and both expression levels are correlated in breast cancer cell lines [57]. *STAT3* and *TNFRSF1A* were also coexpressed in OIR mice (Fig. 7). During angiogenesis and metastasis, there is extensive matrix remodeling, including the production of basement membrane glycoproteins, important for the support and growth of blood vessels. Among them, collagen type IV and laminin are the major constituents of the membrane basement [55,61,62]. Genes for the production of both glycoproteins, *LAMB1* and *COL4A1*, are part of our signature and regulatory RNA network, and both are associated with tumor growth, proliferation, metastasis, and drug resistance in sereval types of cancer [61–67].

Also in the regulatory network we found the *PROS1* gene, which encodes Protein S, a vitamin K-dependent anticoagulant protein, secreted by activated T-cells or tumor-associated macrophages [68,69]. *PROS1* may have antagonist roles as reported by some studies. This gene acts as a tumor suppressor gene in breast and lung cancers. Its overexpression is associated with reduced cell migration and invasion in breast cancer [68,70], and reduced metastasis of lung cancer [68]. However, *PROS1* promotes tumor growth and survival in glioma [69], glioblastoma [71], prostate cancers [72], and squamous cell carcinoma [73]. Our analysis shows that *PROS1* mRNA level was significantly higher in low-risk *HER2*+ breast cancer patients compared to intermediate and high-risk patients (S5 Fig.), corroborating studies that point to a tumor suppressor role in breast cancer. A similar situation exists with the *PLXND1* gene, which encodes a transmembrane protein receptor for semaphorins and neurophilins, and is involved in the regulation of vascular and axon patterning [74,75]. Similar to *PROS1*, *PLXND1* can act as a tumor promoter or as a tumor suppressor depending on the cancer type [74,76,77]. Using METABRIC data, *PLXND1* mRNA level was associated with poor prognosis (HR 1.48, *p*-value = 0.05), and in the *HER2*+ breast cancer subtype *PLXND1* was significantly more abundant in the high-risk patient group, when compared to intermediate and low-risk patients.

Noteworthy is the position of *STAT3* a well-known and key regulatory gene in physiological and pathological angiogenesis, mediating *VEGFA* and *FGF2* cell activation pathways, as well as matrix metalloproteinases (*MMP*) expression [78–80]. *STAT3* is connected in the network to *RHOJ*, for example, which is a GTPase responsible for signaling cytoskeletal rearrangement in endothelial cells during angiogenesis [81–83]. Both genes were tested in different cancer cells lines, and it was established that both stimulate tumor growth and survival by promoting angiogenesis [79–81,83].

For a long time, non-protein-coding regions were called “junk DNA”; in humans these regions account for about 98% of the genome [84]. Now, we know that these non-coding RNAs have important biological functions, including regulation of the angiogenesis process. One example is the DE lncRNA “32952”, which we identified as a potential new isoform of the lncRNA *MEG3* (S1B Fig.). *MEG3* is a maternally expressed imprinted gene, with multiple isoforms. Interestingly, *MEG3* has been described as a tumor suppressor gene and angiogenesis inhibitor [85,86], and has also been implicated in the pathogenesis of ophthalmologic disease [87] and found to be downregulated in many cancer cell types. When overexpressed, it suppresses tumor growth and angiogenesis in different cancer cells [88–91]. In the OIR model, lncRNA “32952” was strongly inhibited on the angiogenesis pick (P17 expression is 17 times lower in the OIR group compared to the control group), which is consistent with the results of the previous studies just mentioned. Together with the mRNAs, our network revealed lncRNAs (including new lncRNAs) coexpressed with key genes of pathological angiogenesis, suggesting they may be regulating their function. Thus, the gene regulation network of pathological angiogenesis helped us identify central RNAs in this process and potential candidates for development of new therapies for breast cancer or other angiogenesis-dependent diseases.

Among the 10 top RNAs with the highest importance and high influence in the pathological angiogenesis network (Fig. 7), there are two new lncRNAs that were predicted in our pipeline, “61969” and “61677”; they are likely antisense lncRNAs. Indeed, lncRNA “61969” is on the opposite strand of gene *RAB12*, which suggests that this lncRNA regulates the activity of the *RAB12* gene. However, these RNAs do not belong to the same co-expressed module. *RAB12* is a small GTPase that is primarily involved in vesicle transport and trafficking, specifically the transportation of transferrin receptors from the endosome to the lysosome [92,93]. *RAB12* has been associated with radioresistance in HPV+ Cervical Cancer Cells [94]. lncRNA “61677” is on the opposite strand of gene *RFX2*. Considering that both belong to the same module of co-expression, our results present strong evidence that this new lncRNA regulates the *RFX2* gene during pathological angiogenesis. *RFX2* is a transcriptional factor which is key to spermatogenesis [95]. A recent study also observed that *RFX2* is involved in the regulation of tumor angiogenesis in kidney renal clear cell carcinoma [96]. In sum, we presented evidence that these three new lncRNAs, “32952”, “61969”, and “61677” are involved in the regulation of the genes from our signature. They are thus potential targets for the development of new drugs to treat breast cancer patients or other angiogenesis-dependent diseases.

In summary, our study indicates that (metastatic) *HER2*+ breast cancer progression relies on a network of *VEGF*-independent pathways, such as those driven by angiopoietin-2 (encoded by *ANGPT2*) and platelet-derived growth factor (*PDGFB*), for tumor vascularization. Of course, it does not rule out the participation of *VEGF*, but if *VEGF* is not available (or inhibited), other *VEGF*-independent pathways may take over and drive tumor growth and progression. One intriguing possibility is vessel co-option. It agrees with the fact that *HER2*+ often metastasize to lung, liver and brain, tissues in which vessel co-option has been well-documented [30,97]. Furthermore, angiopoietin-2, a prominent feature in our gene signature, has been shown to participate and promote this mechanism of tumor vascularization [30,98,99]. Notwithstanding the mechanisms of neovascularization, our novel *HER2*+ gene signature and regulatory RNA network may contribute to researchers in the field seeking alternatives for drugs, diagnosis and therapeutic markers to treat patients with breast cancer.

## Materials and Methods

### Ethics statement

Animal study ethical approval was granted by the Animal Study Ethics Committee from the Institute of Chemistry of the University of São Paulo (approval #10/2010).

### RNA Data

We analyzed RNA-Seq data generated from poly-A enrichment (Poly-A+) and rRNA depletion (Ribo-) libraries constructed using the total RNA extracted from mice retinas of OIR model as previously described [9]. In brief, library preparation and sequencing were performed at the Next Generation Sequencing Core from Scripps Research Institute (San Diego, CA) and strand-specific libraries were enriched for polyadenylated transcripts using the NEB Next Ultra Directional RNA Library Prep Kit for Illumina (New England BioLabs). The retinas of treated (OIR) and non-treated mice were dissected on days 12, 15, and 17, except for the OIR group, for which an extra sample on day 12, called R12.5, was taken, representing the peak of hypoxia in the time series. For the Poly-A+ library, 2 biological replicates and 6 to 7 technical replicates for each biological replicate were performed for each group, except for day 15 when technical replicates were not performed, totaling 69 libraries (S4 Table). For the Ribo-libraries, 2 biological replicates were made for each group for each time, totaling 14 libraries (S5 Table).

### Reference genome

The genome available in public databases (e.g.: ENSEMBL and NCBI) refers to the C57BL/6J strain produced by The Jackson Laboratory. However, the strain used in this study was from the NIH (National Institutes of Health) C57BL/6N, which has some genetic variations compared to the 6J strain [100]. For this, a custom script was developed to modify the reference genome (6J) with the 6N lineage mutations described in Keane et al. [101]. The script is available on GitHub (https://github.com/lbi-usp/rf2m).

### Predicting new lncRNAs

We performed the prediction of new lncRNA transcripts using the sequencing data from the poly-A+ and the Ribo-libraries. To assemble the new transcripts, we used the software StringTie [102], Ryüto [103], and Scallop [104], and merged all assemblies, using the “merge” function of StringTie, into one set of non-redundant transcripts, having as reference the annotation of the mouse genome. We performed a BLAST search (identity ≥ 90%, coverage ≥ 90%, e-value ≤ 10^-5^) against GENCODE (GRCm38.p6) mRNA and ncRNA sequences [105,106] from human, mouse, and rat, and INFERNAL (e-value ≤ 10^-5^) [107] using as reference the covariance models deposited in RFAM [108], to identify whether any of the new transcripts are similar to another lncRNA already described. We also measured the coding potential of the new transcripts with CPAT (coding probability ≤ 0.44) [109] and RNASamba (classification = ‘noncoding’) [110]. A transcript was considered as potential lncRNA if both CPAT and RNASamba were concordant, or if BLAST or INFERNAL returned a valid hit.

### Transcript abundance

RSEM [111] was chosen to quantify the abundance of the transcripts, using as reference the principal isoform of each gene, according to the APPRIS database [112], or the longest isoform if the gene is missing in APPRIS database, plus the lncRNAs predicted according to the methods described in the Methods section.

### Building the co-expressed genes network of angiogenesis

We ran a principal component analysis (PCA) with all samples to check the quality of the replicates (S6 Fig.). We identified a batch effect in the Ribo-library, introduced probably by sequencing the biological replicates in different lanes in the Illumina platform. For this reason, we decided to only use the abundance estimations of the Poli-A+ library for downstream analysis. Transcript expression data were normalized with DESeq2 [113] and those with variance lower than the median (considering the estimated variance in all samples) were filtered out. The co-expressed genes modules were determined using WGCNA [12] (*networkType* = “signed”, *corType* = “bicor”, *corOptions* = list(maxPOutliers = 0.05), *soft threshold power* = 20). The soft threshold power (β) was determined using the WGCNA *pickSoftThreshold* function, where the lowest value capable of generating a network with a scale-free topology (R^2^ = 0.9) was chosen [114]. The genes were clustered using topological overlap measures [12] and the modules were defined using the Dynamic Tree Cut algorithm [115]. Once the modules were identified, we calculated the correlation of the eigengene of the module with each group of samples. Modules with a significant correlation (*p*-value ≤ 0.05) were selected for downstream analysis.

### Building gene signatures for breast cancer

We downloaded the METABRIC dataset [14,15] from the cBioPortal database (https://www.cbioportal.org/), which contains survival data and metadata of ∼2000 breast cancer patients, which is a satisfactory number of samples to train and validate gene signatures. We divided this dataset into 2 cohorts, training and validation (Table 3) with no overlap between them. Univariate Cox regression analysis was conducted with each gene (only those with human homologs) of the selected modules to verify which genes have a significant association (*p*-value ≤ 0.05) with the survival time of the patients from the METABRIC training cohort. For those, we selected the top 50 genes with the highest difference in normalized expression between retinopathy samples and control samples. To build the best gene signature, we ranked the selected genes based on their *p*-values from the Univariate Cox regression analysis and fed the Random Forest algorithm (randomforestSRC R package) [116] with the top gene of the ranked list, always incrementing the signature with the next top ranked gene in each step until all the 50 selected genes were incremented in one of the signatures. The best signature is the one with the minimum error (out-of-bag error).

### Building the mRNA-lncRNA-miRNA network of the gene signature

We recover the protein-protein interaction network of the genes from the Brown signature using STRING database [27], filtering interactions with a minimum confidence score of 0.400 (medium confidence) and excluding those which the interaction source were from text mining. To complement this network, we selected the co-expressed pairs of RNAs from the Brown module of the WGCNA analysis. To restrict the number of interactions for better visualization and analysis of the network, we calculate the 90^th^ percentile of the interaction weights between mRNA-mRNA of genes of the signature, which was 0.2519518, and use this value as a threshold to select the co-expressed pairs of mRNA-mRNA and lncRNA-mRNA with interaction weight above this threshold. We calculated the degree and the betweenness centrality of each node of the resulted network using Cytoscape and created two subnetworks: 1) the 10 RNAs with the highest degree and, 2) the 10 RNAs with the highest betweenness centrality. We incremented these two subnetworks with miRNAs that have the potential to target those RNAs. We searched in the OncomiR [117] database for miRNAs related to cancer development, progression, and survival. To restrict the number of potential miRNA target interactions (MTIs), we verify if the MTIs were previously validated using the miRTarBase database [118], a curated database that contains only MTIs experimentally validated.

## Data availability Statement

RNA-seq data from Poli-A+ mRNA is available in the Sequence Read Archive (SRA) under accession code BioProject PRJNA483866. RNA-seq data for total RNA depleted for ribosomal RNA has been deposited to SRA under accession PRJNA1347474. It will become public upon acceptance.

## Acknowledgements

We thank professor Paul Boutros from the Department of Human Genetics and Urology at the David Geffen School of Medicine at UCLA for helpful technical comments.

## Supporting information

**S1 Fig. Genomic coordinates of the new lncRNAs.** A) 62942, located in the Mtpap gene, and B) 32952, located in the *MEG3* gene.

**S2 Fig. Kaplan-Meier analysis of the Brown and Ivory gene signatures in the LumA, LumB and Basal breast cancer subtypes of the METABRIC cohort.** The p-value was calculated using log-rank test.

**S3 Fig. Genomic coordinates of the new lncRNAs.** A) 61969, which is in the opposite strand of the gene *RAB12*, and B) 61677, which is in the opposite strand of the gene *RFX2*.

**S4 Fig. Kruskal-Wallis test of the differential expression of *A2M* gene in *HER2+* breast cancer patients.** There is a significant difference in the expression of the gene *A2M* in HER2+ breast cancer patients in the Low risk group compared to Intermediate risk group.

**S5 Fig. Kruskal-Wallis test of the differential expression of *PROS1* gene in *HER2+* breast cancer patients.** There are significant differences in the expression of the gene *PROS1* in HER2+ breast cancer patients in the Low risk group compared to Intermediate risk group and High risk group.

**S6 Fig. PCA plot with Ribo- and Poly-A+ library samples.** It is possible to see a clear separation of the Ribo- and Poly-A+ samples concerning the principal component 1. It is also possible to observe segregation between biological replicates 1 and 2 of the Ribo-library, which points to a batch effect presented by sequencing these samples on different Illumina lanes.

**S1 Table. LncRNAs differentially expressed in the OIR retina considering all time points.**

**S2 Table. Genes from each individual module that were combined with patient age and tumor stage to feed the random forest package.**

**S3 Table. Gene signatures that could significantly segregate (*log*-rank test *p*-value ≤ 0.05) cancer patients using the METABRIC validation cohort.**

**S4 Table. Total number of sequenced reads in Poli-A+ library samples.**

**S5 Table. Total number of sequenced reads in Ribo-library samples.**

**S1 File. Genes significantly associated with breast cancer patients’ overall survival (Cox regression univariate analysis p-value ≤ 0.05).**

